# Cell-cycle dependent organization and dynamics of RNA Polymerase I in live human cells

**DOI:** 10.1101/133082

**Authors:** William Conway, Won-Ki Cho, Namrata Jayanth, Susan Mullen, Ibrahim I Cissé

**Affiliations:** Department of Physics, MIT, Cambridge, MA

## Abstract

RNA Polymerase I (Pol I) is responsible for over 60% of transcriptional output in human cells, yet basic questions concerning the spatial and temporal organization of the polymerase remain unanswered. Here we investigate how mammalian cells rely on Pol I organization throughout the cell cycle to balance different needs, from complete transcription shut down to massive increase in protein synthesis (and thus ribosomal RNA synthesis) before cell division. In contrast to our previous reports on RNA Polymerase II, Pol I clusters are stable with active transcription, and the presence of transient Pol I clusters correlates with inactive ribosomal transcription. Our results suggest that both stable and transient populations Pol I clusters co-exist in individual living cells, and their relative fraction may directly reflect the global gene expression need of the cell.

## Introduction

The organization and dynamics of molecular processes inside the cell’s nucleolus, the sub-nuclear domain dedicated to ribosomal biosynthesis, are emerging as a key intersection point for many cellular functions. To balance the demand for ribosomes in cellular growth with the energetic cost of ribosome biosynthesis, cells are believed to have evolved complex regulatory mechanisms to modulate ribosomal synthesis in response to the availability of nutrients and energy (Kusnadi et al. 2015; Schmit 1999). These regulatory mechanisms exhibit significant conservation and cross-talk with pathways critical in the cellular response to environmental stress, cell growth and maintenance, due to the importance of ribosome production in these cellular functions (Zhao et al. 2016; Grummt 2013; Boulon et al. 2010). A majority of regulatory schemes may affect the activity of RNA Polymerase I (Pol I), the molecular complex responsible for the production of ribosomal RNA. Similarly, Pol I is said to be a promising target for the treatment of tumors (Schneider 2012; Quin et al. 2014). Yet, despite the central position of Pol I regulation in coordination of cellular processes, it remains unclear how Pol I organization and dynamics depend on, or reflect, cellular demand for ribosomal RNA transcription.

Classical models of Pol I organization have emphasized the importance of ribosomal (rDNA) gene clustering. There are roughly three hundred copies of the 5.8 kb rDNA gene organized in tandem arrays throughout the human genome (McStay and Grummt 2008). Early electron micrographs revealed that these rDNA genes cluster together within the nucleolus (Miller and Beatty 1969). On each cluster of ribosomal genes, many polymerases simultaneously transcribe pre-rRNA transcripts. When spread for electron microscopy, the nascent pre-rRNAs spread out from the rDNA gene to form a Christmas-tree like, stable structures colloquially termed “Miller spread” (Miller and Beatty 1969).

In the nucleolus of living cell’s during interphase, however, individual subunits of the Pol I holoenzyme exhibit dynamic turnover: different subunits of Pol I display different fluorescence recovery after photobleaching (FRAP) kinetics suggesting that the recruitment of Pol I machinery relies on metastable intermediates (Dundr 2002). Kinetic modeling on FRAP data suggested that there may be multiple subpopulations of Pol I interacting with the ribosomal genes, including one subpopulation with rapid recovery. In fact, studies have suggested that ribosomal transcriptional regulation occurs via cell-cycle dependent modulation of these kinetics by altering the availability of the transcription factor SL1, thereby changing the assembly efficiency of the ribosomal transcription complex (Gorski 2008).

However ensemble-averaging techniques like FRAP are not readily amenable to direct visualization of co-existing subpopulations, and further characterization of how each subpopulation putatively changes to help balance cellular progression through different cycles. Furthermore, previous live cell studies of Pol I relied on the over-expression of components of the polymerase machinery which may alter relative protein abundance. It is unclear whether such over-expression systems accurately reflect endogenous polymerase organization and dynamics.

Here, we employ CRISPR/Cas9 to label endogenous protein, and quantitative super resolution imaging to study the organization and dynamics of endogenous Pol I in live human cells. We fused the catalytic subunit of Pol I, the endogenous RPA190 to a photoconvertible fluorescent protein, Dendra2 (Chudakov et al. 2007), using the CRISPR/Cas-9 genome editing system in a human osteosarcoma (U2OS) cell line (Ran et. al 2013; Jinek et. al 2013; Cong et al. 2013; Mali et al. 2013; Cho et al. 2016b). Upon illumination with 405 nm light, Dendra2 is photoconverted from green to red (Chudakov et al. 2007) allowing for imaging of single RPA190 molecules in living cells. Using this Pol I labeled cell line, we employed quantitative super-resolution imaging to investigate Pol I organization and dynamics in living cells (Cisse et al. 2013; Cho et al. 2016).

## Results and Discussion

To label endogenous Pol I, a homologous donor template containing the Dendra2 insertion (without a stop codon) between a left homologous arm containing 500bp immediately upstream of the gene and a right homologous arm containing the first 500bp of the gene was transfected into U2OS cells (Figure 1a and Table S1 and S2). Through a guide RNA, the CRISPR/Cas-9 system was targeted to induce a nick in the genomic DNA. This nick triggered homologous recombination between the genomic DNA and the homologous donor template resulting in an endogenous Dendra2 tag on the N-terminus of RPA190. Successful integration was confirmed by comparison with an overexpression system (Figure S1) and via FACS sorting (Figure S2).

**Figure 1:**
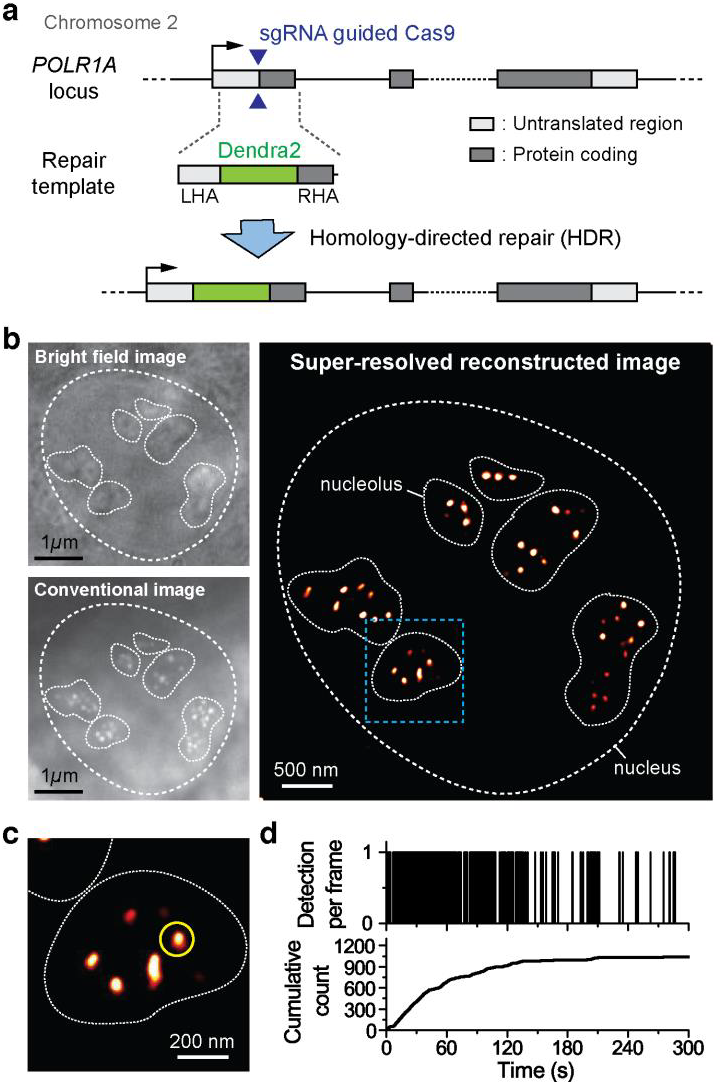
Interphase Organization and Dynamics of RNA Pol I. a) A homologous donor vector containing the Dendra2 fluorescence protein flanked by two 500bp homologous arms. When co-transfected with a plasmid expressing Cas9 along with a targeted sgRNA, homology directed repair at the PAM cut site induces insertion of the Dendra2 sequence onto the N-terminus of the RPA190, the largest subunit of RNA Pol I.b) Bright field and conventional fluorescence imaging of the pre-converted state of Dendra2-RPA190 in a U2OS cell line. The nucleus is demarcated with a dashed line while the contours of the nucleoli are demarcated with a solid white line. The polymerase appears to cluster in distinct foci within the nucleoli. c) Super-resolution reconstruction of the Dendra2-RPA190. At the super-resolution level, foci remain visible. d) A sample time trace of the cluster marked in yellow in the super-resolution image. In the cumulant, the detections show an initial linear slope indicating that the cluster was pre-existing and slowly level off suggesting that the cluster is stable.

With this endogenously labeled Pol I cell line, we used time correlated photoactivated localization microscopy (tcPALM) to investigate the real time, super-resolution dynamics of Pol I (Cisse et al. 2013, Cho et al. 2016a). In single-molecule localization based super-resolution techniques such as (f)PALM or STORM a final image is reconstructed based on precise localizations of a sparse, stochastically activated set of fluorophores (Betzig et al. 2006; Hess et al. 2006; Rust et al. 2006). Time-correlated super-resolution analysis extracts protein dynamics from a super-resolution reconstruction. Since the number of localization events in a temporal window can serve as a measure of local concentration, super-resolution acquisition in live cells can provide a measure of the relative spatial and temporal fluctuations of the labeled proteins.

### RNA Pol I Dynamics in the Interphase Nucleus

We began by investigating the real-time dynamics of Pol I during interphase. Pol I is sequestered in distinct foci within the nucleoli (Figure 1b and 1c) (Dundr et al. 2002). Examining the time traces of the super-resolution localizations using tcPALM reveals that these foci are stable (Figure 1d). The cumulative traces display a steady initial stream of localizations, indicating that the cluster existed before the beginning of image acquisition. The gradual plateau in the tcPALM time trace suggest that the cluster is still present and the pool of available fluorophores is gradually depleted during the imaging process. Consistent with the tcPALM signature for stable clusters, some of the pre-converted Dendra2 Pol I foci also appear stable by direct (conventional) imaging (Figure 1b; lower panel). The presence of multiple stable, sub-nucleolar interphase Pol I clusters is consistent with a picture of the nucleolus sub-organized into distinct regions including dense transcriptional centers (with high Pol I concentration) and outer nucleolar surroundings. During mitosis, tandem arrays of the rDNA genes segregate into regions known as nucleolar organizer regions (NORs) which are responsible for the formation of nucleoli after cell division. Active transcription is thought to occur at the interface of dense fibrillar component and the fibrillar center (Olson and Dundr 2015), and as ribosomes are assembled, they are believed to progress outwards for splicing, processing and association with core ribosomal proteins (Thomson et al. 2013) which putatively make up the rest of the nucleolus surrounding Pol I clusters. Our observation of stable clusters of RNA Pol I within the nucleoli also agree with previous observations of stable ribosomal transcription loci in living mammalian cells (Dundr 2002).

### SL1 Inhibition with CX-5461

Next, we sought to investigate Pol I dynamics at low levels of transcription levels via drug inhibition. We inhibited RNA Pol I initiation with a small molecule inhibitor CX-5461. CX-5461 is a synthetic compound that selectively inhibits RNA Pol I while leaving other transcription (for example mRNA synthesis by RNA Polymerase II) unaffected (Drygin et al. 2011; Haddach et al. 2012). RNA Pol I transcription depends on a variety of transcription factors, chiefly UBF and SL1 (Grummt 2003). UBF binds to the rDNA promoter and recruits SL1. SL1 then recruits the downstream transcription factors and associated machinery needed to bind RNA Pol 1 to the promoter. CX-5461 is thought to act selectively on Pol 1 by inhibiting SL1, thereby preventing RNA Pol I initiation. The drug has further been observed to induce autophagy and prevent cell growth and division. We incubated the cells in a 2μM CX-5461 for 48 hours before performing super-resolution experiments.

The initiation-inhibited cells also show foci of accumulated Pol I, though these apparent foci appear less bright against the background compared to the control, untreated cells by conventional imaging (Figure S2). The foci become much readily visible after super-resolution reconstruction (Figure 2a). The time traces of individual foci, are qualitatively different from the untreated case. In contrast to the gradual plateau of localizations in the untreated foci, many of the initiation-inhibited foci display a transient signature (Figure 2b). Initially in the time trace there are virtually no localizations, suggesting that there was no pre-existing cluster before the start of acquisition. Localization frequency then suddenly increases, indicating that the local concentration of Pol I has rapidly increased in the foci. The localization detections then cease abruptly suggesting that the cluster has likely disassembled. The number of detections in this transient jump results from multiple molecules and can not be accounted for by single molecule photophysics (Cho et al. 2016a). Together, the data illustrated in the example time trace in Figure 2b suggest that the Pol I cluster transiently formed and disassembled upon transcription inhibition, in contrast to the stable Pol I clusters normally observed in interphase in Figure 1d.

**Figure 2:**
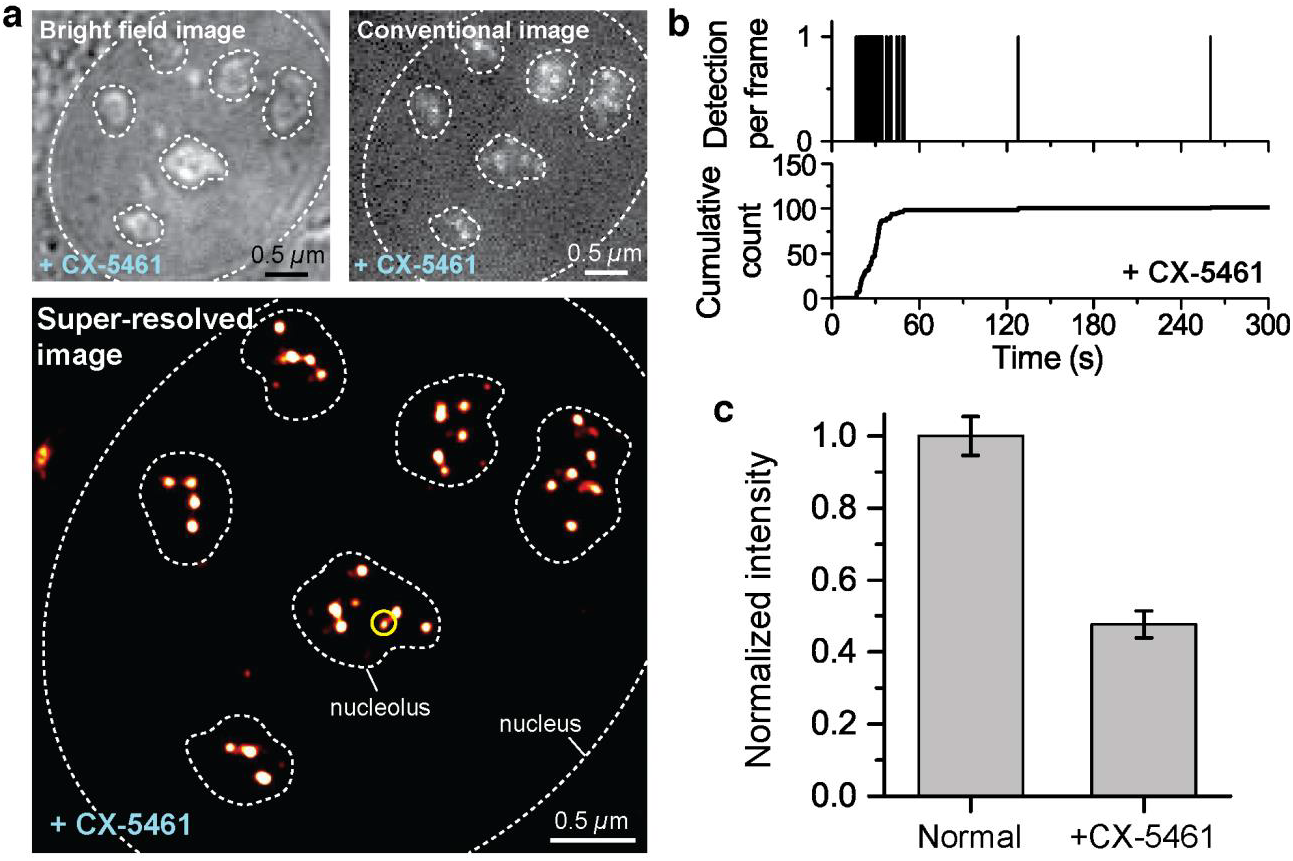
Pol I Reponse to CX-5461 Initiation Inhibition. a) Brightfield, conventional and super-resolution reconstrution images of the Dendra2-RPA190 U2OS cell line after 48 hours of treatment with CX-5461. Conventional flouresence imaging was achieved via 50ms of exposure with a 488nm laser. Sparse foci appear against a stronger background like in the untreated, interphase nucleus. Many dimmer foci that were not visible in the conventional image appear in the super-resolution reconstruction. b) A sample time trace of a transient cluster in the CX-5461 treated nucleus. c) Raw intensities of pre-converted Dendra2-Pol I in normally grown cells and CX-5461-treated cells were measured. N=55 foci from 16 normally grown cells and N=45 foci from 13 CX-5461-treated cells were collected. 200 frames of images were averaged for each cell. Pol I foci of the CX-5461 were less than half as bright as the foci of normally grown cells (*I*=0.48 ± 0.04; mean ± s.e. of mean).

The onset of transient clustering yields insights into the role of polymerase in the stable clusters. Inhibiting initiation affects cluster stability suggesting that stable clusters of Pol I are attributable to elongating polymerases on the rDNA gene. When the polymerase is unable to bind, we see a transient accumulation of polymerases rather than the stable signature. Some stable clusters persisted after treatment, likely due to the inefficiency of the drug in turning off all ribosomal transcription.

### Cell-Cycle Dependence of RNA Pol I Dynamics

To explore polymerase organization at varying transcription levels, we characterized the cell cycle dependence of RNA Polymerase I organization. We investigated the dynamics of Pol I in mitosis. We blocked cells in S phase by halting the cell cycle using a double thymidine block (Bostock et al. 1971 and see Methods and Materials). Releasing these cells from thymidine produces synchronized cells in M and G1 phases (Bostock et al. 1971 and see Methods and Materials).

The RNA Pol I spatial organization in M phase differs substantially from interphase. We observe no groups of distinct foci of polymerase clusters in M phase, likely because the chromosomes are condensed. In a select few cells (∼10% of cells imaged), there is a small number of polymerase foci visible (Figure 3a). In the M phase cells, we observe a mixture of both stable and transient Pol I foci (Figure S3).

**Figure 3:**
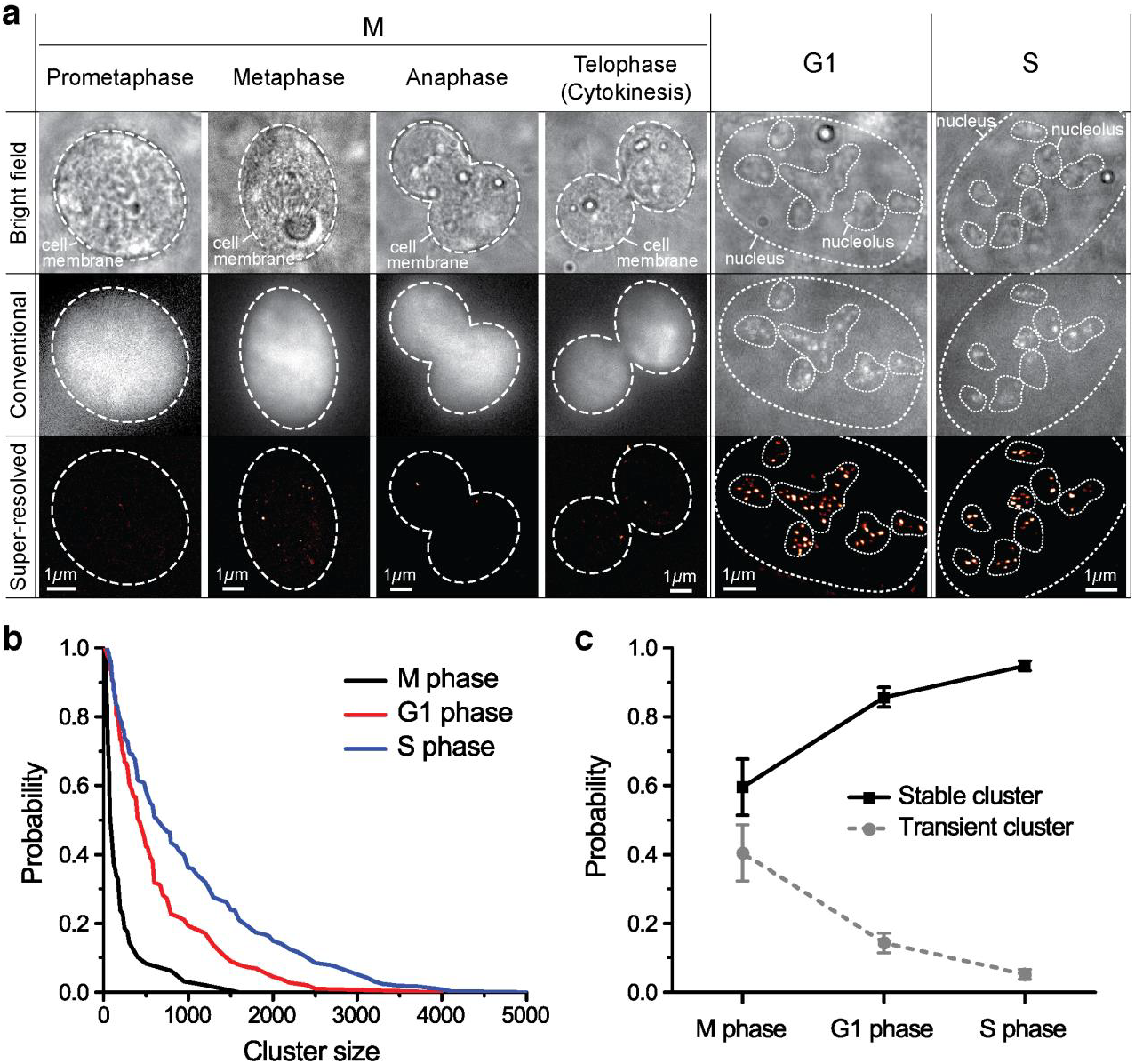
Cell-cycle dependent organization and dynamics of RNA Polymerase I. a) Bright field, conventional and super-resolution images of M, G1 and S phase cells. b) Survival curve of stable cluster size in M, G1 and S phase cells. c) Portion of stable and transient clusters in M, G1 and S phase cells.

During mitosis, the chromosomes condense and transcription of rDNA ceases (Klein 1999). Previously bound Pol I molecules are thought to remain bound to the rDNA genes during this transcriptionally silent period (Roussel 1996). Novel Pol I binding events, however, are believed to be inhibited, putatively by Runx2 which forms a complex with the UBF and SL1 transcription factors, inhibiting initiation (Young et al. 2007). Our observations agree well with our expectations from induced ribosomal transcription inhibition. Though we observe fewer clusters in mitosis than in inhibited interphase, we see a mix of transient and stable clusters similar to that observed with transcription inhibition. The stable clusters are attributable to polymerases that putatively remain bound to the rDNA genes during mitosis while the transient clusters linked to the absence of the SL1 as we observed when we inhibited SL1 with CX-5461(Roussel 1996; Grummt 2003; Young et al. 2007).

To quantify the differences in Pol I foci dynamics between different phases of the cell cycle, we evaluated both the portion of transient and of stable clusters and the size distribution of the stable clusters (Figure 3b and 3c). We found that M phase displays the highest fraction of transient clusters, and the fraction of transient clusters decreases as cells progress into G1 and later S phase.

Similarly, we quantified the relative intensity of stable clusters. We observe more detections per stable clusters as cells progress out of M phase into G1 and S (Figure 3b). Translating number of detections into number of molecules is an intricate task in super-resolution data due to non-trivial single-molecule photophysics. Nonetheless, more detections likely indicates that more polymerase are present in the foci. Thus, Pol I clusters tend to become more intense (i.e. larger by the number of molecules per foci) and more stable as the cells grow and synthesize new DNA in preparation for division.

We note that the observation of larger, more stable clusters corroborates previous interpretations from FRAP data showing longer retention times for individual polymerase subunits and the SL1 transcription factor in S phase than G1 (Gorski et al. 2008). The increase in Pol I cluster stability and size also corroborates well with previous observations that Pol I activity increases as cells progress through G1 and peaks in S phase (Grummt et al. 2013).

However, the simultaneous presence of a transient population, distinct from stable foci could not be inferred in previous bulk measurements. The fraction of stable versus transient clusters may be interpretable as a relative fraction actively transcribing rDNA versus inactive, silent nucleolar organizer regions, a measure that may likely reflect the overall activity of the rDNA genes.

Taken together, our live cell super-resolution data paint a physical picture of cell cycle dependent RNA polymerase I activity and organization (Figure 4) whereby Pol I forms stable foci in the nucleolus where rDNA genes are clustered and when many polymerases are actively transcribing the rDNA genes. When polymerase cannot bind to the promoter, Pol I may still cluster albeit very transiently. Transcriptionally active nucleolar centers thus appear as stable clusters of polymerase while inactive nucleolar centers appear as transient clusters. This is in sharp contrast to RNA Polymerase II in the nucleoplasm, where transient clusters form during initiation of actively transcribed genes (Cho et al. 2016a, Cisse et al. 2013), and clusters are stabilized with drugs that inhibit phosphorylation and prevent promoter escape.

**Figure 4:**
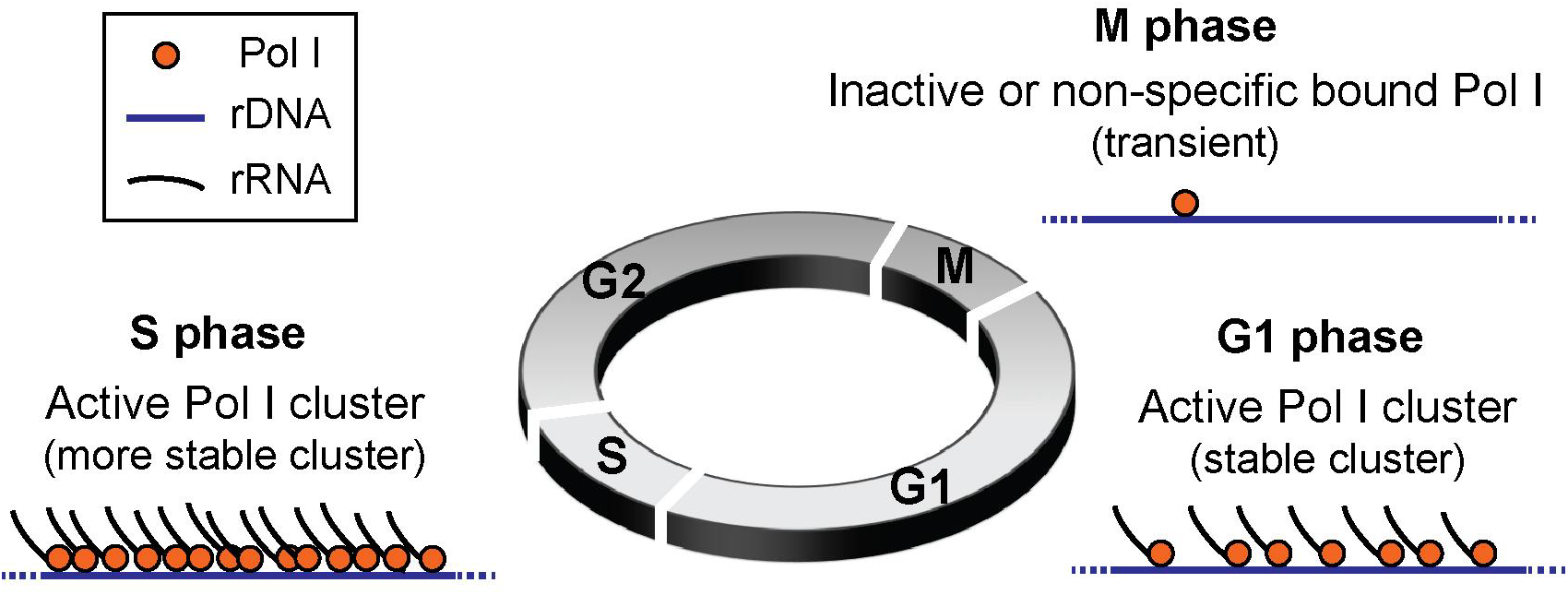
Cell-cycle dependent organization and dynamics RNA Polymerase I. During M phase, we observe inactive transient clusters of RNA Polymerase I. As cells recover from division and progress towards DNA replication in S phase, active stable clusters of RNA Polymerase I form. These clusters consist of many polymerase bound to each ribosomal gene actively transcribing pre-rRNA. The size of clusters grows as cells progress from G1 to S phase indicating more bound transcribing polymerases.

## Methods and Materials

### CRISPR-Cas9 mediated insertion of Dendra2 onto the N-Terminus of RPA190

A human osteosarcoma (U2OS) cell line with an endogenous N-terminal Dendra2 insertion in the RPA190 gene was generated via the CRISPR/Cas9 system (Cong et al. 2013).

The cells were transfected with both a px459 plasmid containing an sgRNA targeted to start of the RPA190 gene (5’-TTCAGCCGAATACATCCCCGAAGG-3’) and a homology directed repair template. sgRNA sequences were generated via the online CRISPR toolbox (crispr.mit.edu) and then cloned into the px459 vector with one step digestion ligation (Table S1) (Hsu et al., 2013). All experiments and validation were performed on the cell line transfected with sgRNA1. Of the remaining guides, only sgRNA5 showed successful insertion.

The homologous donor sequence was synthesized by Life Technologies to contain a 500bp left homologous arm, the 690bp Dendra2 insertion and a 500bp right homologous arm. Silent mutations were introduced into the PAM sites of 6 potential sgRNAs in the right homologous arm to ensure the repair template was not degraded by the Cas9 system. The repair template was PCR amplified to prepare a linear fragment for transfection.

The linear homologous repair template and the px459 plasmid containing the sgRNA insert were transfected into the U2OS cell line using the xTreme9 transfection reagent from Sigma-Aldrich. Cells were left to incubate at 37°C for 24 hours. The cells were then allowed to recover, and sorted via FACS at the Koch Institute Sorting facility to isolate cells expressing the Dendra2 insertion for imaging.

### Cell Culture Protocols

The Dendra2-Pol I cells were cultured in Dulbecco’s Modified Eagle Medium with Glutamax (DMEM with Glutamax) from Thermo Fisher (10567). The media contained 10% fetal bovine serum from Glibco (26140-079, US Origin, Qualified) and an antibiotic mixture at a final working concentration of 10U/mL penicillin and 10 μg/ml streptomycin from Gibco (15140). Cells were incubated at 37° C with 5% supplemental CO2.

### Initiation Inhibition with CX-5461

Cells were incubated with 2μM CX-5461 (Selleckchem 1138549-36-6). CX-5461 is a potent, selective inhibitor of SL1, a transcription factor associated with Pol I binding to the rDNA gene (Drygin et al. 2011; Haddach et al. 2012). CX-5461 was stored at 10mM in a stock solution of 50mM NaH2PO4 (pH 4.5) and added directly to imaging dishes for treatment.

### Cell Cycle Synchronization via a Double Thymidine Block

Cells were synchronized using a double thymidine block approach previously reported (Bostock et al. 1971). Cells were treated with 2mM of thymidine from Sigma Aldrich (T1895-1G) dissolved in the 10% FBS DMEM media previously described and incubated at 37° C for 15 hours to arrest cells in S phase. Following this initial S phase arrest, cells were released via removal of the thymidine medium and allowed to progress for 9 hours to ensure all cells had passed out of S phase. Cells were then reincubated with 2mM thymidine to synchronize cells at the G1/S junction. To produce cells in S phase, cells were imaged ∼one hour after release from thymidine control. For M phase, cells were imaged between 9 and 12 hours after release from thymidine and subject to visual confirmation of ongoing mitosis (i.e rounded cell shape, finger-like projections, cell division, etc.). For G1 cells, cells were imaged 15 hours after thymidine release.

### Super-Resolution Imaging

Cells were imaged on a homebuilt super-resolution setup, comprised of a Nikon Eclipse TI microscope equipped with a 100x oil immersion objective (NA 1.40) (Nikon, Tokyo, Japan) and lasers and filter sets for 405nm, 488nm, and 561nm illumination. During imaging, cells were kept at 37° C in an incubator set atop the objective (InVivo Scientific, St. Louis, MO). Images were captured on an Andor iXon Ultra 897 EMCCD camera at a rate of 60ms/frame at an EM-gain of 900. Camera image acquisition and control was performed using Micro Manager 1.4 (Edelstein et al. 2014). Cells were held steady in the z-direction using the Perfect Focus System of the Nikon Microscope during image acquisition.

For PALM imaging, excitation (561nm) and activation (405nm) lasers were combined, expanded then focused on the sample. These beams were expanded using an achromatic beam expander (AC254-040-A and AC508-300-A, from THORLABS, Newton, NJ) and refocusing was performed with an achromatic converging lens (#45–354, from Edmund Optics, Barrington, NJ). Power levels were controlled both by directly varying the initial laser intensity and through an AOTF, and measured directly at the top of the objective lens closest to the sample slide.

### Image Analysis and tcPALM

We analyzed images following the general scheme previously described in depth in Cho et al. 2016 and Cisse et al. 2013. Individual Pol I molecules were localized using a modified MTT localization algorithm (Sergé et al. 2008). Super-resolution images were generated from a Gaussian spreading of the localizations determined by the MTT program. We then analyzed these localizations using a homebuilt, open source software (qSR) for analysis of both spatial and temporal correlation of localization events available for free on the Cisse lab’s Github page (www.github.com/cisselab/qSR) (Andrews J.O. et al. in preparation).

## Acknowledgements

We would like to thank Cisse Lab members Arjun Narayanan, Takuma Inoue, and Jan-Hendrick Spille (MIT) for helpful comments, and J Owen Andrews for comments and assistance with analysis software. Research reported in this publication was supported by the National Cancer Institute of the National Institutes of Health under the NIH Director’s New Innovator Award (DP2CA195769) to I.I.C. The content is solely the responsibility of the authors and does not necessarily represent the official views of the National Institutes of Health. This work was also supported by funds from the MIT Department of Physics, the DeFlorez Endowment Fund, and the MIT Undergraduate Research Opportunities Program (UROP).

